# Teaching the Skills and Concepts of Gene Expression Analysis during COVID-19

**DOI:** 10.1101/2022.06.28.497969

**Authors:** Louis A. Roberts, Scarlet S. Shell

## Abstract

Teaching students how to conduct impactful gene expression studies within an authentic research paradigm typically relies on hands-on experience to develop skills at the bench, coupled with conceptual understanding of the experiments and their analyses within or outside of the lab setting. The unexpected shift of our gene expression laboratory course to a remote model provided the opportunity to assess how student learning of these skills and concepts would be affected in an entirely online environment. Our data suggest the reduced learning gains for skill-based techniques came with increased student learning of concepts and analysis, which may have aided stimulating student interest in gene expression studies.

---

Our department offers a novel upper-level “Molecular Biology and Genetic Engineering: Approaches and Applications” laboratory course based on the program of one of our research-active faculty members. Based on the research of Dr. Scarlet Shell (Associate Professor, BBT), students clone a DNA sequence (e.g., the 5’UTR of a ribosomal protein gene) into a shuttle vector with a standard promoter designed to express Yellow Fluorescent Protein (YFP). Students integrate their reporter constructs into *Mycobacterium smegmatis* (a non-pathogenic model for *M. tuberculosis*) and stress the cells with ribosome-targeting antibiotics. We hypothesize 5’ UTRs may alter reporter transcript/protein levels upon stress as measured by quantitative PCR (qPCR), plate reader, and flow cytometry. This ambitious experimentation takes place over seven weeks; most students complete the methods and acquire data, but time for detailed analyses is limited.

When COVID-19 necessitated teaching the entire lab remotely, we sought to at a minimum retain the typical level of conceptual understanding of the experimental workflow, methods, and analyses, and acknowledged the lack of bench work would impair the skills-based components of the course. We wondered if time lost in lab could be reallocated to increase student learning of how data are analyzed and presented, both from individual experiments and within peer-reviewed journals. We used data obtained from previous offerings of the course, we benefited from having data generated by several course alums who continued their projects in the Shell Lab (one of whom co-authored a peer-reviewed journal article[1]). We utilized two instruments-university student course reports, and a “Skills and Concepts Inventory” (SCI) we created-to gauge student learning by comparing the online (2020) and in-person (2019) versions of the lab.

The SCI asks students to self-rate their knowledge/comfort with a given skill or concept they encounter in the course (0-4 scale), and is administered on the first and last days of lab to calculate learning gain (LG). The SCI data show the in-person version rated higher at primarily skills-centric learning (e.g., sterile technique, micropipetting and assays, microscopy, PCR, cloning, mRNA/cDNA methods; see Figure 1). The university student course reports (1-5 scale, completed at the conclusion of the course; see Table 1) also are notably higher when students are assessing skills development in the in-person modality.

**Table 1.** University Student Course Report Data.

| # | Question text | In person (n=8) | Remote (n=11) |
| --- | --- | --- | --- |
| Q1 | My overall rating of the quality of this course is | 4.3 | 4.1 |
| Q3 | The educational value of the assigned work was | 4.1 | 4.4 |
| Q4 | The instructor's organization of the course was | 3.9 | 3.9 |
| Q7 | Relative to other college courses I have taken, the amount I learned from the course was | 4.5 | 4.3 |
| Q8 | Relative to other college courses I have taken, the intellectual challenge presented by the course was | 4.0 | 4.1 |
| Q10 | The instructor stimulated interest in the subject matter | 4.0 | 4.7 |
| L1 | The instructor showed me how to use lab equipment properly | 4.3 | 3.8 |
| L2 | The lab and/or computer equipment was in good operating condition | 4.5 | 3.6 |
| L3 | Good lab practices were emphasized | 4.5 | 4.6 |
| L4 | Relative to other lab experiences, the intellectual challenge presented by the lab assignments was | 3.9 | 4.3 |
| L5 | Relative to other lab experiences, the clarity and specificity of lab assignment objectives was | 4.1 | 3.9 |

n= number of student responses.

**Figure 1.**
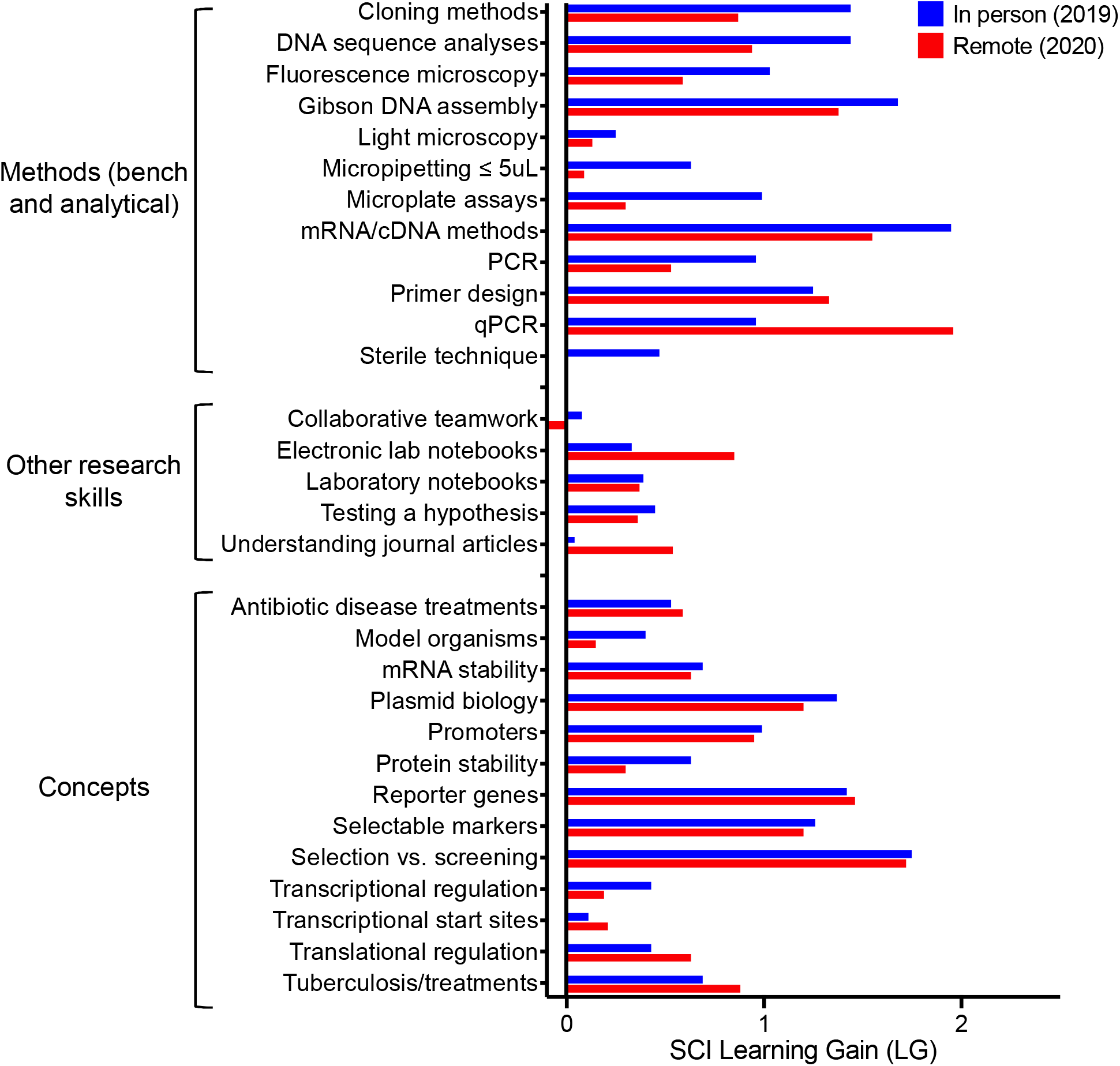
The learning gains measured via the Skills and Concepts Inventory (eight student responses for in person version, and 11 for remote). See S1 for SCI instrument.

Conversely, both the SCI and the university course report data strongly suggest student conceptual understanding was similar in both versions of the lab. University course report data show the overall course rating and perceived quantity and value of what was learned were comparable between offerings. Many of the concepts that underpin the experiments (i.e., plasmids, selection, screening, reporters, primer design, lab notebooking) have similar SCI LGs for each version, as do the motivations for performing the research (i.e., tuberculosis disease treatment and antibiotic resistance mechanisms). Surprisingly, the key concepts of qPCR, construction of journal articles, and electronic lab notebooking showed dramatic LG increases in the online version according to SCI data. qPCR differs from regular PCR primarily in that it produces a different data type requiring conceptually complex analysis. We also note that the highest increase in the university course report data (4.7 online vs. 4.0 in-person) was in response to the degree to which the course “stimulated interest in the subject matter”.

In summary, the apparent increase in conceptual understanding by the students, coupled with the course inspiring student interest in molecular biology applications, indicate the potential strengths and value of instructional methods and platforms that transcend teaching skills at the lab bench.

## Supporting information

Supplemental SCI survey instrument

**S1:** Skills and Concepts Inventory (SCI) survey instrument.

## Notes

### Competing Interest Statement

The authors have declared no competing interest.

