## Supplemental SCI survey instrument for "Teaching the Skills and Concepts of Gene Expression Analysis during COVID-19"

**Skills and Concepts Inventory (SCI)**

Please complete the following questionnaire regarding your familiarity with the following components of our course. You will be asked to complete this survey at the conclusion of the term.

Please rank your familiarity/comfort/experience with the following on a 0-4 scale, where…

0 is “I completely unfamiliar/have no experience”

1 is “I have heard of this in theory/done this once but am not comfortable”

2 is “I am somewhat comfortable with this”

3 is “I am familiar with this concept/technique”

4 is “I am an expert with this concept/technique”

**Sterile/aseptic technique with any living cell type**

0	1	2	3	4

**Micropipetting volumes ≤ 5uL**

0	1	2	3	4

**Light microscopy**

0	1	2	3	4

**Fluorescence microscopy**

0	1	2	3	4

**Microplate activity/detection assays (i.e., enzymes, fluorescent molecules, ELISAs, Bradford/A660/BCA)**

0	1	2	3	4

**Assess validity of a hypothesis via experimental design, data collection, and analysis**

0	1	2	3	4

**Keeping a proper laboratory notebook**

0	1	2	3	4

**Electronic laboratory notebooking (Benchling, Lab Archives, etc.)**

0	1	2	3	4

**Peer-reviewed journal article structure, format, and content**

0	1	2	3	4

**Performing science as a collaborative team**

0	1	2	3	4

**Polymerase chain reaction**

0	1	2	3	4

**Designing primers to amplify/modify a double-stranded DNA template**

0	1	2	3	4

**Cloning methods**

0	1	2	3	4

**DNA assembly to build plasmids (HiFi, Gibson, etc.)**

0	1	2	3	4

**Annotation and alignments of DNA sequences**

0	1	2	3	4

**mRNA isolation and cDNA synthesis**

0	1	2	3	4

**Quantitative PCR (qPCR, RT-PCR)**

0	1	2	3	4

**Plasmid biology**

0	1	2	3	4

**Selectable markers and selection**

0	1	2	3	4

**How selection differs from screening (in the context of transformation)**

0	1	2	3	4

**Promoters**

0	1	2	3	4

**Definition of transcriptional start sites**

0	1	2	3	4

**Reporter genes**

0	1	2	3	4

**Antibiotic treatment of disease**

0	1	2	3	4

**Tuberculosis infection and treatment**

0	1	2	3	4

**Model organisms in science**

0	1	2	3	4

**Transcription-level gene regulation**

0	1	2	3	4

**Translation-level gene regulation**

0	1	2	3	4

**mRNA stability**

0	1	2	3	4

**Protein stability**

0	1	2	3	4
